# Scene perception and visuospatial memory converge at the anterior edge of visually-responsive cortex

**DOI:** 10.1101/2022.11.14.516446

**Authors:** Adam Steel, Brenda D. Garcia, Kala Goyal, Anna Mynick, Caroline E. Robertson

## Abstract

To fluidly engage with the world, our brains must simultaneously represent both the scene in front of us and our memory of the immediate surrounding environment (i.e., local visuospatial context). How does the brain’s functional architecture enable sensory and mnemonic representations to closely interface, while also avoiding sensory-mnemonic interference? Here, we asked this question using first-person, head-mounted virtual reality (VR) and fMRI. Using VR, human participants of both sexes learned a set of immersive, real-world visuospatial environments in which we systematically manipulated the extent of visuospatial context associated with a scene image in memory across three learning conditions, spanning from a single field-of-view to a city street. We used individualized, within-subject fMRI to determine which brain areas support memory of the visuospatial context associated with a scene during recall (Exp. 1) and recognition (Exp. 2). Across the whole brain, activity in three patches of cortex was modulated by the amount of known visuospatial context, each located immediately anterior to one of the three scene perception areas of high-level visual cortex. Individual subject analyses revealed that these anterior patches corresponded to three functionally-defined place memory areas, which selectively respond when visually recalling personally familiar places. In addition to showing activity levels that were modulated by the amount of visuospatial context, multivariate analyses showed that these anterior areas represented the identity of the specific environment being recalled. Together, these results suggest a convergence zone for scene perception and memory of the local visuospatial context at the anterior edge of high-level visual cortex.

**Significance statement:** As we move through the world, the visual scene around us is integrated with our memory of the wider visuospatial context. Here, we sought to understand how the functional architecture of the brain enables coexisting representations of the current visual scene and memory of the surrounding environment. Using a combination of immersive virtual reality and fMRI, we show that memory of visuospatial context outside the current field-of-view is represented in a distinct set of brain areas immediately anterior and adjacent to the perceptually-oriented scene-selective areas of high-level visual cortex. This functional architecture would allow efficient interaction between immediately adjacent mnemonic and perceptual areas, while also minimizing mnemonic-perceptual interference.

## Introduction

Perception is shaped by our immediate sensory input and our memories formed through prior experience (Albright, 2012; Rust and Palmer, 2021). Yet, how memory influences ongoing perception in the brain is poorly understood. A particular puzzle is how the brain enables sensory and mnemonic representations to functionally interface, while avoiding interference that occurs when the same neural population concurrently represents both sensory and mnemonic information (Rust and Palmer, 2021). One elegant system for studying these questions is scene perception, which relies heavily on concurrent mnemonic-sensory interactions: as we explore our environment, memory of the visuospatial content immediately outside of the current field-of-view informs what we will see next as we shift our gaze (Haskins et al., 2020; Draschkow et al., 2022).

How does the brain’s functional architecture support coexisting representations of the current visual scene and memory of the surrounding environment? Recent studies suggest that scene-related mnemonic and sensory information converge at the anterior edge of high-level visual cortex (Baldassano et al., 2013, 2016; Silson et al., 2016; Bainbridge et al., 2020; Popham et al., 2021; Steel et al., 2021). For example, we recently localized brain regions that preferentially activated when participants recall personally familiar real-world places (e.g., their house) versus faces (e.g., their mother). This revealed three patches of cortex located anterior and adjacent to one of the three classic “scene perception areas” of high-level visual cortex that selectively activated when recalling familiar places (Steel et al., 2021). These “place memory areas” constituted a distinct functional network from the scene perception areas and co-fluctuated with both spatial memory structures (i.e., the hippocampus) and scene perception areas, bridging mnemonic and perceptual systems. Based on these results, we proposed that these paired perceptually- and mnemonically-oriented functional networks may, respectively, represent scenes and their remembered visuospatial context. This possibility is attractive because it would allow efficient interaction between adjacent mnemonic and perceptual areas while minimizing mnemonic-perceptual interference (Steel et al., 2021).

However, while previous work identified place-recall driven activity anterior to each of the scene-perception areas, the content of the representations in these anterior regions is currently unclear. This is partly because prior studies investigating place memory, including our own, used participants’ personally familiar real-world places as stimuli (e.g., ‘your mother’s kitchen’) (Peer et al., 2015, 2019; Steel et al., 2021). While personal memories elicit strong responses and are appropriate for some experimental questions, these memories contain rich associations that go beyond visuospatial content (i.e., the appearance of your mother’s kitchen), including semantic knowledge (e.g., where her kitchen is located), multisensory details (e.g., the smell of baking cake), and episodic context (e.g., blowing our birthday candles). As a result, it is unclear whether visuospatial memory *per se* is sufficient to engage the anterior memory areas. Likewise, whether the anterior areas process information about the visuospatial context of a scene is currently unknown.

To address this knowledge gap, we used head-mounted virtual reality (VR) to teach participants a set of real-world environments. This approach allowed us 1) to ensure participants’ memories contained no confounding episodic information and limited semantic content, and 2) to precisely manipulate the visuospatial context associated with each environment. We defined visuospatial context as the spatial extent, configuration, and visual details of the local environment. Next, we used individualized, within-subject fMRI to test whether, across the brain, the place memory areas play a special role in processing visuospatial memory. Specifically, we asked whether these areas represent the extent of the visuospatial context associated with a visual scene during recall (Exp. 1) and visual recognition (Exp. 2) of single scene views from each place (Figure 1B-C). We hypothesized that if the place memory areas process visuospatial context associated with a scene view, i) their activity will reflect the degree of associated visuospatial context (Bar and Aminoff, 2003; Baumann and Mattingley, 2016) and ii) they should contain multivariate representations of the scene’s specific visuospatial content.

**Figure 1.**
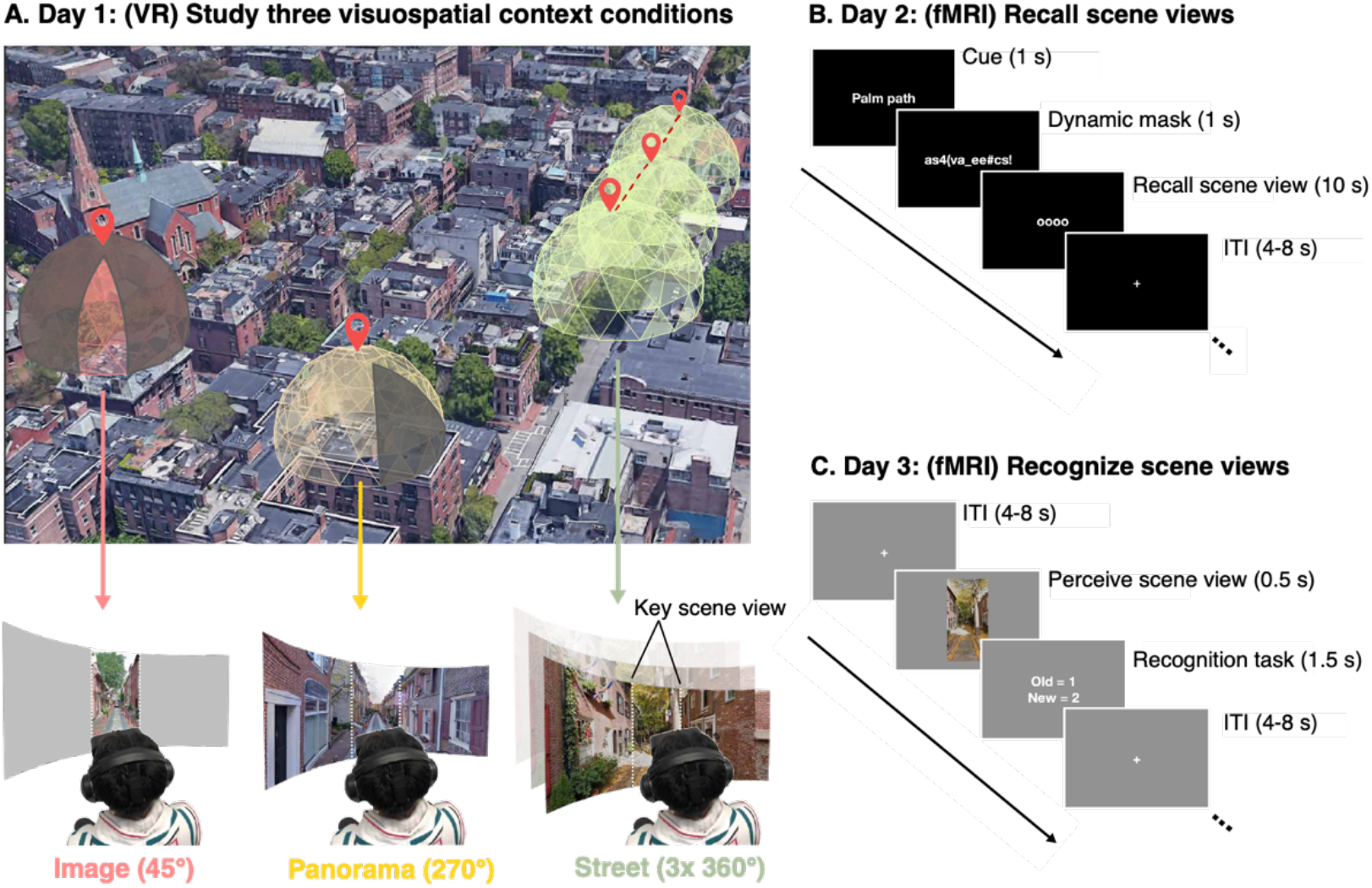
Experimental design to test processing of visuospatial context associated with a real-world scene. The experiment took place over across three sessions, one study session, and two fMRI sessions. These sessions took place on three separate days. A. In the VR study session, participants studied 20 real-world visual scenes with varying amounts of visuospatial context in head-mounted virtual reality (VR). Each scene was associated with one of three levels of spatial visuospatial context: single images (45° visible, 315° occluded), panoramas (270° visible, 90° occluded), and streets (three contiguous, navigable 360° photospheres). During the two fMRI sessions, we tested the neural response during recall (B) and recognition (C) of the learned scenes using fMRI.

## Methods

### Participants

17 subjects (10 Female; age=21.6 ± 2.9 std years) participated in this experiment. This number was based on a power calculation from our prior work, specifically based on the average effect size observed when participants viewed panning movies of familiar versus unfamiliar scenes across ROIs (Cohen’s D = .76 with 80% power and Alpha = 0.05 requires a minimum of 16 subjects; Experiment 2, (Steel et al., 2021)). Specifically, from our prior work, we calculated the difference in activation between the place memory and scene perception areas on the lateral, ventral, and medial surfaces to familiar versus unfamiliar videos, as this was our best proxy for visuospatial context. We then took the average difference score across cortical surfaces for each subject and used these values for our power calculation. We felt confident assuming this large effect size because of the consistency in the effect across subjects (all subjects showed greater familiarity effects in the place memory areas compared to the scene perception areas).

Participants were Dartmouth students and postdocs recruited via posters and word-of-mouth advertising. They received monetary compensation for taking part in the study. Participants had normal or corrected-to-normal vision, were not colorblind, and were free from neurological or psychiatric conditions. Written consent was obtained from all participants in accordance with the Declaration of Helsinki with a protocol approved by the Dartmouth College Committee for the Protection of Human Subjects (CPHS).

### Procedure

The study took place over three sessions (each on a different day): one study session and two fMRI sessions. During the study session, participants studied 20 real-world places using a head-mounted virtual reality (VR) system (see Stimulus Presentation section). The 20 places comprised three spatial context conditions (see Stimuli section). During the first fMRI session (Experiment 1), we investigated which brain areas represent the level of known spatial context by assessing the neural response across the three spatial context conditions using fMRI. During the second fMRI session (Experiment 2), we tested whether the same regions that represent context during recall also represent known spatial context during visual recognition by evaluating neural responses when viewing images of the studied places. 17 participants took part in Experiment 1, and 16 participants took part in Experiment 2.

### Stimuli – Study Phase

Stimuli consisted of 20, 360° panoramic photospheres of outdoor real-world places from 5 cities (Oslo, Philadelphia, Padua, Quebec, Seville), sourced from Google Maps using iStreetView (istreetview.com). Participants were not told about the real-world locations from which the photospheres were taken. Stimuli can be found on Open Science Framework in the ‘panoramas’ folder: https://osf.io/3ygqb/?view_only=5f436411c021448abf5157df3f2b2f30. At the end of the second fMRI session, we asked participants whether they recognized any photospheres from real-life (i.e., had they been to any of these locations outside of VR). Importantly, no participant recognized any stimuli, suggesting that real-world familiarity could not explain our results.

Example videos from the study phase on Open Science Framework in the ‘trial videos’ folder: https://osf.io/3ygqb/?view_only=5f436411c021448abf5157df3f2b2f30. Stimuli were applied to a virtual environment created in Unity (www.unity3d.com), and the experimental routine was programmed using custom scripts written in C#.

The photospheres were divided into three spatial context conditions: 1) single images (45° from a photosphere, 315° occluded by a grey screen), 2) panoramas (270° visible, 90° occluded), and 3) streets (three contiguous 360° photospheres tiling a city street, navigable via button press). The cities were represented equally in all conditions (i.e., from each city, 2 photospheres were used in as image trials, 1 photosphere was a panorama trial, and 3 contiguous photospheres comprised a street trial). During the study session, we occluded a portion of the photosphere in the Image and Panorama conditions to obscure the wider visuospatial context. Note that, because of the unique demands of curating and presenting photospheres for the street condition, all photospheres were assigned to the same conditions across participants. Therefore, it is possible that different levels of memory for the photospheres could contribute to a Main effect of condition in Exps. 1-2. Importantly, however, our *a priori* hypotheses deal with interactions between conditions and ROI (i.e., differences in activation magnitude between perceptual and mnemonic areas), which would not be impacted by this main effect of condition. Moreover, we found no differences across conditions in our behavioral assessments of memory, suggesting that condition-related differences we observed in neural activity were not driven by stimulus-related effects.

Unbeknownst to the participants, the image condition was comprised of two sets of five images derived from i) the occluded portion of the panorama condition and ii) an isolated photosphere from an independent location within the same city. We found no difference between the image conditions in any regions of interest during the recall and recognition experiments (Experiment 1 (recall): OPA: t(16)=0.95, p=0.35; PPA: t(16)=0.81, p=0.43; MPA: t(16)=0.58, p=0.56; LPMA: t(16)=0.83, p=0.42; VPMA: t(16)=0.84, p=0.41; MPMA: t(16)=0.75; p=0.46; Experiment 2 (recognition): OPA: t=1.84, p=0.086; PPA: t(15)=0.0007, p=.99; MPA: t(15)=0.75, p=0.46; LPMA: t(15)=0.013, p=0.99; VPMA: t(15)=0.86, p=0.40; MPMA: t(15)=0.63, p=0.53), so we considered these images together as a single Image condition.

The Street condition consisted of three contiguous photospheres representing a city street. After entering the trial, participants rotated 180° from their initial facing direction to see an arrow on the floor, which indicated that they could advance forward. This rotation ensured that the key scene view presented during fMRI (i.e., initial viewpoint) did not depict the extra visuospatial context experienced by the participant. Participants advanced forward by pressing a button on a handheld controller. Participants could advance forward two positions, allowing them to view around a street corner unobservable from the initial position. Participants could not go backward to revisit photospheres during the street trial (i.e., they could only travel ‘one-way’).

### Stimulus presentation

During the study phase, experimental stimuli were presented to participants via an immersive, head-mounted virtual reality display (16 Oculus Quest 2, 1 Oculus Go; Fast-switch LED display panel 1832×1920 resolution per eye; 98° field of view; 120 Hz refresh rate). Participants stood wearing the head-mounted VR display during the experiment. This setup offered participants a self-directed opportunity to explore the naturalistic environment from an egocentric perspective naturally via eye-movements and head turns. All fMRI stimuli were presented using Psychopy 3 (Version 3.2.3) (Peirce, 2007).

### Procedure - Study phase

During the study session, participants studied all 20 scenes (20s each). Study trials automatically terminated after 20 seconds. On each trial, participants were instructed to look naturally around each environment, just as they would in real life. Importantly, although the view was restricted in the Image and Panorama conditions, participants could still turn their heads and move their eyes to naturally explore the environment. Therefore, the conditions only differed in the amount of spatial context that participants learned.

Before and after each study trial, participants saw an associated place “name” (e.g., “Scallop path”) that later served as a cue during the fMRI recall experiment. We instructed participants to learn these name-place pairings for a later memory test. After each experimental trial, participants returned to a virtual home screen (a platform surrounded by clouds), where they were instructed to take a break. Time between trials was self-paced (participants initiated the beginning of a new trial via a button press).

During the study session, participants saw the 20 scenes grouped in blocks. No scene was repeated during a study block. Scene order was randomized in each study block. All participants completed at least 5 study blocks (approximately 1 hour of training, depending on subjective confidence and availability). After the 5^th^ block, we asked participants if they were confident that they could recall each place; if participants were not confident, they completed one additional study block. Importantly, all analyses below are within-subject, minimizing the potential impact of across-subject differences in study session duration or stimulus familiarity.

All participants completed a “refresh” study block immediately before entering the scanner on both scan sessions.

### FMRI

#### Experiment 1: Memory recall

The goal of Experiment 1 was to determine what brain areas represent the known visuospatial context associated with a scene view. To this end, we compared BOLD activation when participants recalled places that varied in their degree of known spatial context.

Experiment 1 was performed across eight runs. Each place stimulus was cued once in each run (20 trials/run). On each trial, participants saw the name of a place from the study phase (1 s), followed by a dynamic mask of random alphanumeric characters that rapidly changed (1 s). After the mask, the recall cue (‘oooo’) appeared on the screen for 10 s. We instructed participants to perform mental imagery of a single field-of-view from the prompted place in as much detail as possible while the recall cue was on the screen. The specific instructions given to the participant are reproduced below.

> “During scanning you will perform the memory recall task. In this task, you will see the name of a place that you studied, followed by some scrambled letters and numbers. Then four circles will appear on the screen. When the four circles are on the screen, we want you to visually recall a view from the cued place as vividly as you can. When you are doing the imagery, you should only imagine a single image, whatever the first image that comes to mind when you are cued for that place. So, for Church Street, you might immediately recall the church at the top of the hill - you should hold that image in your mind the whole time and not try to look around the place. When the cue goes away, a cross will appear. When the cross is on the screen, clear your mind and prepare for the next trial.”

Imagery trials lasted 10 seconds and were separated by a jittered interstimulus interval (range 4-8 s). The stimulus order was pseudo-randomized within each run so that conditions could only repeat 2 times consecutively in an imaging run. Although stimuli were not presented in an optimized sequence, we confirmed that our regressors were minimally collinear (correlation between regressors < 0.2 for 100% of task regressors across participants) post-hoc using afni’s 1d_tool.py.

#### Experiment 2: Spatial context during recognition

In Experiment 2, we assessed whether the same regions that represent known spatial context during recall also represent contextual information during visual recognition. For this experiment, we compared neural activity when participants viewed briefly presented images while they performed an orthogonal familiarity judgement task. We hypothesized that by having subjects perform a task that was unrelated to spatial context, any activation that was modulated by spatial context would reflect implicit reactivation of contextual knowledge.

Experiment 2 was comprised of eight runs. Participants performed a familiar/novel judgement task. On each trial, participants saw a single 45° portion of a photosphere (500 ms), and then indicated whether they had studied the image in VR (‘familiar’) or not (‘novel’) (1500 ms). Trials were separated by a jittered interstimulus interval (range 4-8 s). Each familiar stimulus was presented once in each run, along with four novel stimuli (24 trials/run). The stimulus order was pseudo-randomized so that conditions could only repeat two times per run. For consistency, all stimuli were images looking down the center of a street, which limited the amount of identifying visual information the participant saw. Because overall accuracy was high (greater than 92% correct), all trials were included in the analysis.

Novel stimuli for each run were randomly sampled from a set of 10 images participants had not experienced during the study session (2 novel images per city). One image was taken from the unseen portion of the photosphere from the panorama condition (180° from the studied view). The other was taken from a photosphere that had not been experienced by the participant and always comprised a view looking down the street. The photospheres were chosen to be visually similar to the other stimuli from their respective city.

Although stimuli were not presented in an optimized sequence, we confirmed that our stimulus time series were minimally collinear (collinearity < 0.2 for 99.9% of task regressors across participants) post-hoc using afni’s 1d_tool.py

#### Localizer tasks

In addition to the main experimental tasks, participants underwent localizers to establish individualized regions of interest for the scene perception areas (PPA, OPA, MPA) and place memory areas (VPMA, LPMA, MPMA). We used the same localization procedure defined in our prior work (Steel et al., 2021), briefly described below.

##### Scene perception localizer

To localize the scene perception areas, participants passively viewed blocks of scene, face, and object images presented in rapid succession (500 ms stimulus, 500 ms ISI). Blocks were 24 s long, and each run consisted of 12 blocks (4 blocks/condition). There was no interval between blocks. Participants performed two runs of the scene perception localizer.

##### Place memory localizer

The place memory areas are defined as regions that selectively activate when an individual recalls personally familiar places (i.e., their kitchen) compared with personally familiar people (i.e., their mother). To establish individualized stimuli, prior to fMRI scanning, we had participants generate a list of 36 personally familiar people and places (72 stimuli total). These stimuli were generated based on the following instructions.

> “For your scan, you will be asked to visualize people and places that are personally familiar to you. So, we need you to provide these lists for us. For personally familiar people, please choose people that you know in real life (no celebrities) that you can visualize in great detail. You do not need to be in contact with these people now, as long as you knew them personally and remember what they look like. So, you could choose a childhood friend even if you are no longer in touch with this person. Likewise, for personally familiar places, please list places that you have been to and can richly visualize. You should choose places that are personally relevant to you, so you should avoid choosing places that you have only been to one time. You should not choose famous places where you have never been. You can choose places that span your whole life, so you could do your current kitchen, as well as the kitchen from your childhood home.”

During fMRI scanning, participants recalled these people and places. On each trial, participants saw the name of a person or place and recalled them in as much detail as possible for the duration that the name appeared on the screen (10 s, no dynamic mask). Trials were separated by a variable ISI (4-8 s). Place memory areas were localized by contrasting activity when participants recalled personally familiar places compared with people (see ROI definitions section).

#### MRI Acquisition

All data was collected at Dartmouth College on a Siemens Prisma 3T scanner (Siemens, Erlangen, Germany) equipped with a 32-Channel head coil. Images were transformed from dicom to nifti format using dcm2niix (v1.0. 20190902) (Li et al., 2016).

##### T1 image

For registration purposes, on Day 1, a high-resolution T1-weighted magnetization-prepared rapid acquisition gradient echo (MPRAGE) imaging sequence was acquired (TR = 2300 ms, TE = 2.32 ms, inversion time = 933 ms, Flip angle = 8°, FOV = 256 × 256 mm, slices = 255, voxel size = 1 ×1 × 1 mm). T1 images were segmented and surfaces were generated using Freesurfer (Dale et al., 1999; Fischl et al., 2002)(version 6.0) and aligned to the fMRI data using align_epi_anat.py and @SUMA_AlignToExperiment (Argall et al., 2006; Saad and Reynolds, 2012).

On Day 2, we collected a fast anatomical scan (MPRAGE; TR = 2300 ms, TE = 2.32 ms, inversion time = 933 ms, Flip angle = 8°, FOV = 256 × 256 mm, slices = 255, voxel size = 1 ×1 × 1 mm, GRAPPA factor=4). The Freesurfer reconstructions from the scan collected in Day 1 were registered to this scan for analysis of Experiment 2 using @SUMA_AlignToExperiment.

##### Functional MRI Acquisition

To mitigate the dropout artifacts and allow advanced BOLD preprocessing, FMRI data were acquired using a multi-echo T2*-weighted sequence (Poser et al., 2006; Kundu et al., 2012, 2017; Posse, 2012). The sequence parameters were: TR=2000 ms, TEs=[14.6, 32.84, 51.08], GRAPPA factor=2, Flip angle=70°, FOV=240 x 192 mm, Matrix size=90 x 72, slices=52, Multi-band factor=2, voxel size=2.7 mm isotropic. The initial two frames of data acquisition were discarded by the scanner to allow the signal to reach steady state.

##### Preprocessing

Multi-echo data processing was implemented based on the multi-echo preprocessing pipeline from afni_proc.py in AFNI (Cox, 1996). Signal outliers in the data were attenuated (3dDespike)(Jo et al., 2013). Motion correction was calculated based on the second echo, and these alignment parameters were applied to all runs.

The data were then denoised using multi-echo ICA denoising (tedana.py (Kundu et al., 2012; Evans et al., 2015; DuPre et al., 2019, 2021)) to improve overall data quality in ventral temporal and parietal cortex (Steel et al., 2022). In brief, the optimal combination of the three echoes was calculated, and the echoes were combined to form a single, optimally weighted time series (T2smap.py, included in the tedana package). PCA was applied, and thermal noise was removed using the Kundu decision tree method. Subsequently, ICA was performed, and the resulting components were classified as signal and noise based on the known properties of the T2* signal decay of the BOLD signal versus noise. Components classified as noise were discarded, and the remaining components were recombined to construct the optimally combined, denoised timeseries.

Following denoising, images were smoothed with a 5 mm gaussian kernel (3dBlurInMask), and signals were normalized to percent signal change.

#### ROI definitions

##### Scene perception and place memory areas (functionally defined)

Scene perception regions were established using the same criterion used in our prior work (Steel et al., 2021). Scene and face areas were drawn based on a general linear test comparing the coefficients of the GLM during scene versus face blocks. These contrast maps were then transferred to the SUMA standard mesh (std.141) using @SUMA_Make_Spec_FS and @Suma_AlignToExperiment (Argall et al., 2006; Saad and Reynolds, 2012). A vertex-wise significance of p < 0.001 along with expected anatomical locations was used to define the regions of interest (Julian et al., 2012; Weiner et al., 2018). As a control region, the FFA was defined using the contrast of scenes versus faces using the same procedure.

To define category-selective memory areas, the familiar people/places memory data was modeled by fitting a gamma function of the trial duration for trials of each condition (people and places) using 3dDeconvolve. Estimated motion parameters were included as additional regressors of no-interest. Polynomial regressors were not included in multi-echo data analysis. Activation maps were then transferred to the SUMA standard mesh (std.141) using @SUMA_Make_Spec_FS and @Suma_AlignToExperiment. People and place-memory areas were drawn based on a general linear test comparing coefficients of the GLM for people and place memory. A vertex-wise significance threshold of p < 0.001 was used to draw ROIs.

To ensure our ROIs were comparable in size, we further restricted all functional ROIs to the top 300 most selective vertices (Steel et al., 2021). Importantly, our results were consistent using 300-vertex ROIs defined using the ROI center of mass (rather than most selective vertices) as well as 600 vertices rather than 300, suggesting that the results did not depend on the ROI definition.

##### Anatomical regions of interest

The hippocampus and primary visual cortex (occipital pole) were defined for each participant using FreeSurfer’s automated segmentation (aparc + aseg).

### FMRI analysis

#### Experiment 1

##### Deconvolution

For Experiment 1, we employed two standard general linear model (GLM) analyses for i) group whole-brain analysis and ii) univariate and multivariate ROI analyses.

For the group-based whole brain analysis, our general linear model included regressors for each condition (Image, Panorama, Street) that considered each trial using a gamma function of duration 10 s. In addition, a regressor for the memory cues and dynamic masks of each trial was included as a regressor of no interest. These regressors, along with motion parameters and 4^th^ order polynomials were fit to the data. Activation maps were then transferred to the SUMA standard mesh (std.141) using @SUMA_Make_Spec_FS and @SUMA_AlignToExperiment (Argall et al., 2006; Saad and Reynolds, 2012).

For ROI univariate and multivariate analyses, we analyzed data using a general linear model fit with each trial modeled individually for each run using 3dDeconvolve. The memory recall period for each trial was modeled as a single regressor using a gamma function of duration 10 s. In addition, a regressor for the memory cue and dynamic mask was included as a regressor of no interest using a gamma function of duration 1 s. These regressors, along with motion parameters and 4^th^ order polynomials were fit to the data. Activation maps were then transferred to the SUMA standard mesh (std.141) using @SUMA_Make_Spec_FS and @SUMA_AlignToExperiment (Argall et al., 2006; Saad and Reynolds, 2012).

##### Whole-brain analysis

To locate where in the brain neural activity during recall was modulated by spatial context, we conducted a whole-brain analysis on the cortical surface. Beta values for each condition from all subjects were compared using a within-subject ANOVA (AFNI 3dANOVA2, type 3) with Condition (Image, Panorama, Street) as a factor and Participant as a random effect. Data were thresholded at a vertex-wise FDR-corrected p-value (q) =0.05 (F=6.4, p<0.0001). The result of this analysis (i.e., Main effect of Condition) was compared to the location of the place memory and scene perception areas from an independent group of participants (Steel et al., 2021).

To determine whether the neural activity of any subcortical or allocortical structures (i.e., hippocampus) was modulated by spatial context, we conducted a volumetric whole-brain analysis. Participants’ anatomical data was registered to standard space (MNI template) using auto_warp.py and the registration parameters were applied to their statistical maps. The beta values for each condition were then compared using a within-subject ANOVA (AFNI 3dANOVA2, type 3) with Condition (Image, Panorama, Street) as a factor and Participant as a random effect. No subcortical or allocortical regions showed an effect of Condition even at a modest threshold (p=0.05, uncorrected).

##### ROI analysis: Univariate

The t-values for all trials in all runs were extracted from all ROIs (scene perception areas [PPA, OPA, MPA], place memory areas [VPMA, LPMA, MPMA], hippocampus, V1, FFA; see section *ROI definition*, below). These t-statistics were averaged across runs and conditions, and these average t-statistics were compared using an ANOVA implemented in Matlab (Matlab, Mathworks, Natick, Massachusetts).

For comparison between the scene perception and place memory areas, ANOVAs were conducted separately for each surface (Ventral, Lateral, Medial) with Condition (Image, Panorama, Street), ROI (Scene perception, Place memory), and Hemisphere (left, right) as factors. For all other regions (Hippocampus, V1, FFA), separate ANOVAs were used, with Condition (Image, Panorama, Street) and Hemisphere (left, right) as factors. There was no significant effect of hemisphere in any test (all ps>0.05), and thus data are presented collapsed across hemisphere. Post-hoc tests were implemented using Bonferroni-corrected paired t-tests.

##### ROI-analysis: Multivariate

We sought to determine whether the scene perception and place memory areas contained representations of the specific places that participants recalled. In other words, we tested whether we could decode stimulus identity during trials of the recall experiment. To this end, we performed an iterative split-half correlation analysis and looked for correspondence in the spatial pattern of neural activity between each place across the two halves of the data. We focused on identity decoding indices because condition (i.e., image, panorama, street) decoding could be impacted by the differing response magnitude.

For this analysis, we iteratively half split our runs, such that we created all possible combinations of half split data (total half splits = 70; iteration 1: 1,2,3,4 v 5,6,7,8; iteration 2, 1,2,3,5 v 4,6,7,8; and so on). For each iteration, we averaged t-values for each place from half of the runs (e.g., odd versus even) together for each voxel within each ROI. We then calculated the correlation matrix (Spearman’s rho) across halves. This yielded a similarity matrix of all images for each ROI. We then calculated the identity decoding index as the difference in correlation between the pattern of response to a place with itself (across independent halves of data) compared to the correlation of the places with the other members of its visuospatial context condition (i.e., images were correlated with images, panoramas with panoramas, and streets with streets). This ensured that condition-related effects could not explain our decoding results. This was iteratively calculated for all combinations of runs.

Fisher-transformed correlation values for the scene perception and place memory areas on each surface were separately compared to hippocampus and control regions (V1 and FFA) using an ANOVA, with ROI (SPA, PMA, Hippocampus, FFA, V1) as a factor and Subject as a random effect. Post-hoc tests were implemented using Bonferroni-corrected paired t-tests.

#### Experiment 2

Data was analyzed using a general linear model fit separately for each run using 3dDeconvolve. Each familiar image trial was modeled as a single regressor using a gamma function of duration 2 s. In addition, a regressor for unfamiliar images was included as a regressor of no interest using a gamma function of duration 2 s. Because unfamiliar images were not trial unique (i.e., some unfamiliar stimuli were repeated across runs) and differed in number from the familiar images, we could not analyze these data. These regressors, along with motion parameters and 3^rd^ order polynomials were fit to the data. Activation maps were then transferred to the SUMA standard mesh (std.141) using @SUMA_Make_Spec_FS and @SUMA_AlignToExperiment.

##### Whole-brain analysis

To locate where in the brain neural activity during recall was modulated by spatial context, we conducted a whole-brain analysis on the cortical surface. Beta values for each condition from all subjects were compared using a within-subject ANOVA (AFNI 3dANOVA2, type 3) with Condition (Image, Panorama, Street) as a factor and Participant as a random effect. Data were thresholded at a vertex-wise FDR-corrected p-value (q) =0.05 (F=8.1, p<0.0001). The result of this analysis (i.e., Main effect of Condition) was compared to the location of the place memory and scene perception areas from an independent group of participants (Steel et al., 2021).

To determine whether the neural activity of any subcortical or allocortical structures (i.e., hippocampus) was modulated by spatial context, we conducted a volumetric whole-brain analysis. Participants’ anatomical data was registered to standard space (MNI template) using auto_warp.py and the registration parameters were applied to their statistical maps. The beta values for each condition were then compared using a within-subject ANOVA (AFNI 3dANOVA2, type 3) with Condition (Image, Panorama, Street) as a factor and Participant as a random effect. As we found in experiment 1, no subcortical or allocortical regions showed an effect of Condition even at a modest threshold (p=0.05, uncorrected).

##### ROI analysis

In Experiment 2, we sought to determine whether the regions in which activation reflected spatial context during recall also reflect this information during visual recognition. So, we employed the same univariate ROI analysis approach taken in Experiment 1.

The t-values for all trials in all runs were extracted from all ROIs (Scene perception areas [PPA, OPA, MPA], Place memory areas [VPMA, LPMA, MPMA], Hippocampus, V1, FFA; see section *ROI definition*, below). These t-statistics were averaged across runs and conditions, and we compared these average values using an ANOVA implemented in MATLAB.

For comparison between the scene perception and place memory areas, ANOVAs were conducted separately for each surface (Ventral, Lateral, Medial) with Condition (Image, Panorama, Street), ROI (Scene perception, Place memory), and Hemisphere (left, right) as factors. For all other regions (Hippocampus, V1, FFA), separate ANOVAs were used, with Condition (Image, Panorama, Street) and Hemisphere (left, right) as factors. There was no significant effect of hemisphere in any test (all ps>05), and thus data are presented collapsed across hemisphere. Post-hoc tests were implemented using Bonferroni-corrected paired t-tests.

### Post-scan questionnaires

#### Vividness

To ensure that any differences in neural responses across the conditions during recall were not due to a difference in vividness of imagery, we queried participants on their experience after the scan. Specifically, participants rated each stimulus on the overall vividness of their visual imagery on a 1-5 scale (1: Not able to visualize, 5: As though they were looking at the image). We compared the average ratings across conditions using an ANOVA with Condition (Image, Panorama, Street) as a Factor and Subject as a random effect.

#### Image naming test

To ensure participants could remember the trained images, participants performed an image naming test after completing Experiment 2. Eleven participants completed the recognition test; the remaining participants were not able to be contacted. One participant performed the naming task after substantial delay (> 2 weeks after fMRI scanning) and was therefore excluded from this analysis. Participants were shown the images used in the Experiment 2 scan session and had to report the place name. We compared the accuracy across the conditions using an ANOVA with Condition (Image, Panorama, Street) as a Factor and Subject as a random effect.

## Results

Participants (N=17) learned a set of 20 immersive, real-world visuospatial environments in head-mounted VR. Each environment depicted a small portion of a city (e.g., Padua, Seville, etc.) where participants had never been. These environments comprised three levels of visuospatial context: 1) single images (45° from a photosphere, 315° occluded by a grey screen), 2) panoramas (270° visible, 90° occluded), and 3) streets (three contiguous 360° photospheres tiling a city street, navigable via button press) (Figure 1A, see Methods). In the street condition, the final photosphere allowed the participant to look around a corner to view visuospatial information that could not be seen from their original position, which ensured that the known visuospatial context in this condition was greater than the panorama condition. The cities were equally balanced across the conditions to control for visual and semantic components unique to each city.

Participants explored each environment from a first-person perspective using head turns and eye movements for 20-seconds and learned: i) each visuospatial environment and ii) a name associated with each environment. After studying, participants could vividly imagine each scene and accurately recall each scene-name pairing (see ‘Post-scan behavioral results confirmed memory success’, below). Then, we used fMRI to assess which brain areas process visuospatial context during recall (Exp. 1; Figure 1B) and visual recognition (Exp. 2, Figure 1C). In these tasks, participants either visually recalled or viewed pictures of a single scene view from each place (the key scene view, Figure 1).

We began by identifying any brain areas where visuospatial context associated with scenes learned in VR modulated recall-related activation using an exploratory whole-brain ANOVA with Condition (Image, Panorama, Street) as a factor and Subject as a random effect. In other words, we tested where in the brain activity was modulated by the extent of visuospatial context associated with a scene. The logic of this test builds on studies in the object memory literature showing that association cortex responds more strongly to objects with strong associated context (e.g., a bowling pin, which is strongly associated with a bowling alley) vs. weak associated context (e.g., a chair) (Bar and Aminoff, 2003; Baumann and Mattingley, 2016). These univariate effects are thought to reflect activation of brain areas that harbor information from memory associated with a percept – in our case visuospatial context associated with a scene.

This group-level, whole-brain analysis revealed several distinct clusters in posterior cerebral cortex that were modulated by known visuospatial context (Figure 2). Three clusters, in particular, were located immediately anterior to the three scene perception areas (OPA, PPA and MPA), as identified in an independent group of participants (Steel et al., 2021). In fact, the location of these clusters closely corresponded to the location of the “place memory areas”, three brain areas on the brain’s lateral (LMPA), ventral (VMPA), and medial surfaces (MPMA) that selectively respond when participants recall or perceive familiar places, as defined by a group mask from our previous work (Steel et al., 2021). On the ventral surface, the cluster fell at the anterior edge of the VPMA, which largely overlaps with the parahippocampal cortex (Glasser et al., 2016), a region known to represent an animal’s relative position within an environment (Julian et al., 2018; LaChance et al., 2019; LaChance and Taube, 2022) (Figure 2). These results confirm that the place memory areas, which we previously showed are activated during recall of personally familiar places (Steel et al., 2021), are active during recall of a purely visuospatial environment (i.e., a photosphere learned in VR, which is devoid of episodic or semantic information). Having found that the place memory areas are engaged during visuospatial recall even when episodic and semantic information is controlled, we went on to test whether these areas play a role in processing visuospatial context.

**Figure 2.**
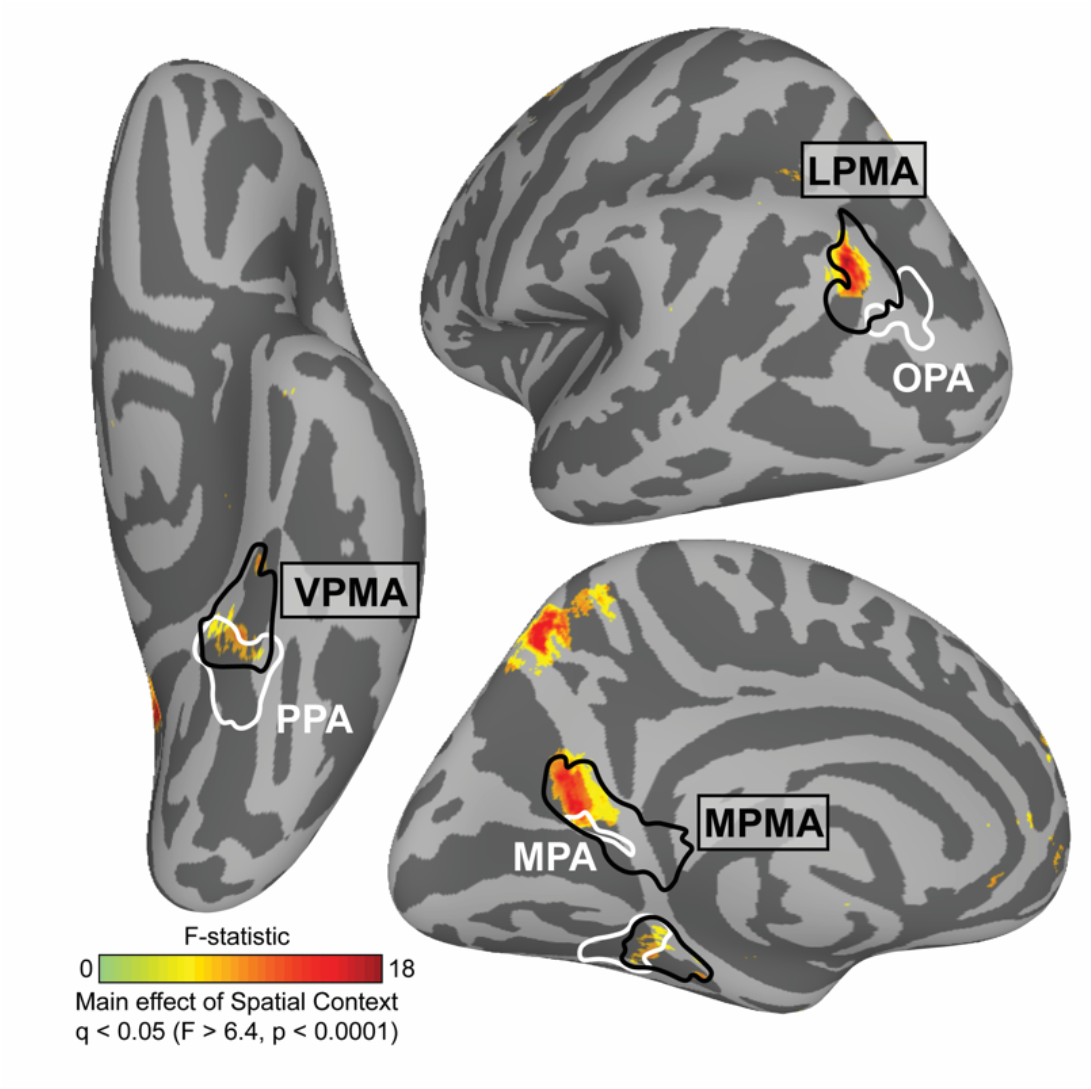
Regions in posterior cerebral cortex are modulated by the extent of known visuospatial context during recall in a group analysis. We compared neural activation across the visuospatial context conditions (Image, Panorama, and Street) using a vertex-wise ANOVA. We found that three clusters showed significant modulation by known spatial context. By comparing these clusters to our prior work mapping perception and memory related responses (Steel et al., 2021), we determined that these regions fell anterior to group-average locations of the scene perception areas (white) and within the place memory areas (black). Whole brain analysis thresholded at FDR-corrected p-value (q) < 0.05 (vertex-wise F>6.4, p<0.0001). Scene perception areas (white) and place memory areas (black) were defined using data from an independent group of participants (Steel et al., 2021).

### Place memory area activity reflects known visuospatial context associated with a scene view during recall

While whole-brain analyses are powerful exploratory tools, they obscure the highly detailed and idiosyncratic location of functional areas present at the individual participant-level (Braga and Buckner, 2017; Gordon et al., 2017; Braga et al., 2019; DiNicola et al., 2020). So, to determine whether the place memory and scene perception areas have dissociable roles when processing visuospatial context, we adopted an individualized ROI-based approach. We independently localized each participant’s OPA, PPA, MPA, and place memory areas on the lateral, ventral, and medial surfaces (LMPA, VPMA, MPMA) (see Methods). We then compared regional activation when participants recalled (Exp. 1) or visually recognized (Exp. 2) the scene views across the three visuospatial context conditions. For each cortical surface, we used an ANOVA with Condition (Image, Panorama, Street), ROI (scene perception/place memory area), and Hemisphere (lh/rh) as factors and Subject as a random effect. Because we found no effect or interaction with hemisphere, results are presented averaged across hemispheres.

### Visuospatial context specifically modulates place memory area activity during recall

This analysis revealed a remarkable dissociation between the scene perception areas and their anterior, memory-driven counterparts (the place memory areas) (Figure 3A). Namely, on all cortical surfaces, visuospatial context modulated place memory area activity, while the scene perception areas were largely unaffected.

**Figure 3.**
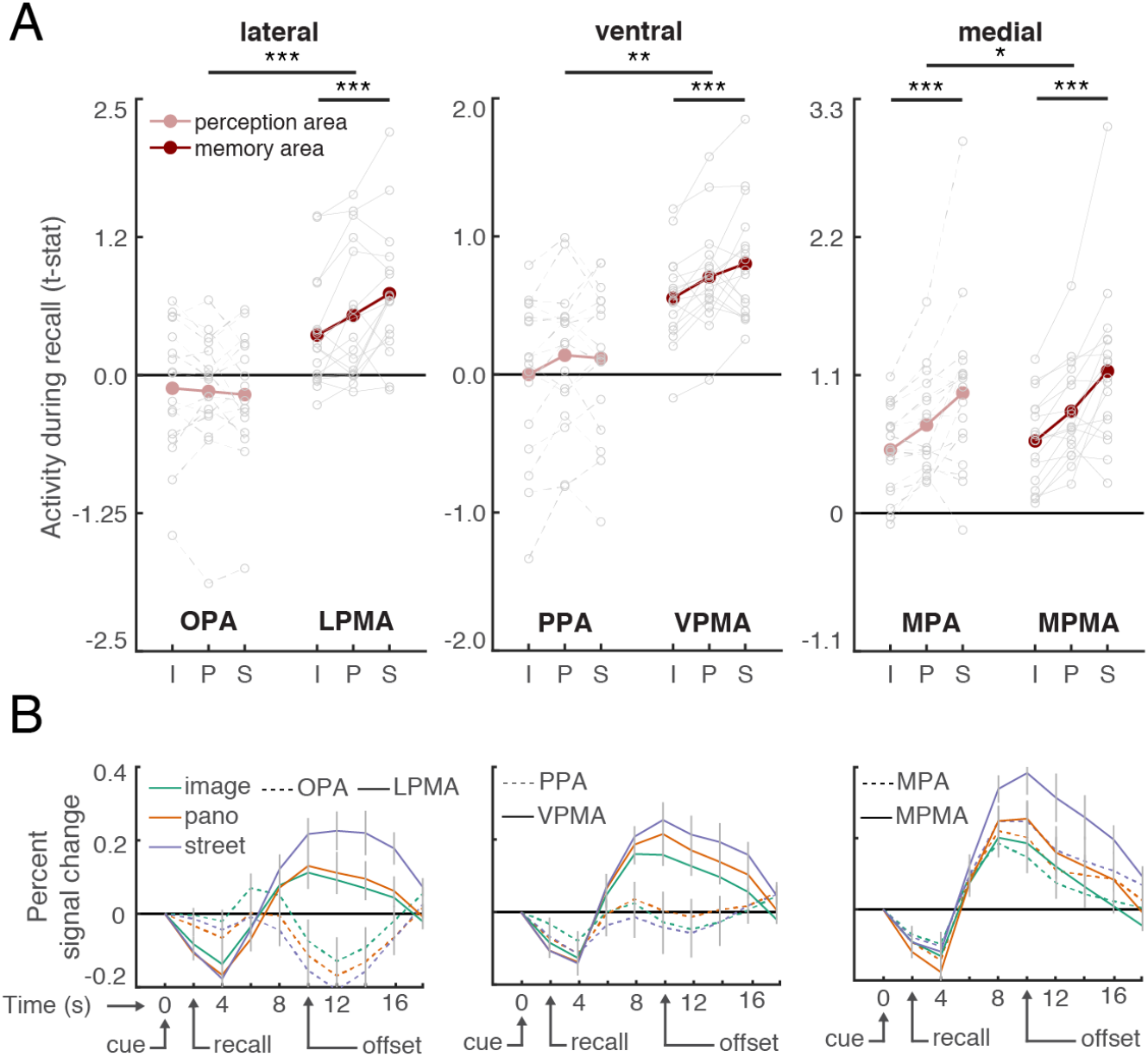
The extent of visuospatial context associated with a scene view modulates place memory area activation during recall. A. For each participant, we compared activation of individually-localized scene perception and place memory areas across the visuospatial context conditions during recall using an ANOVA with ROI (perception/memory area), visuospatial context (image, panorama, street [i,p,s]) and hemisphere as factors. Across all cortical surfaces, the place memory area activation was modulated by spatial context to a greater degree than the scene perception areas. Grey dots indicate individual participants. See Figure S1.1-1.3 for control areas. B. Time course of activation during recall (in seconds) shows differential responses of the scene perception and place memory areas. On all cortical surfaces, the place memory areas showed robust activation during recall (4-12 seconds), while the scene perception areas on the ventral and lateral surfaces did not. Grey lines indicate SEM at each TR (2s). *: p<0.05, **: p<0.01, ***: p<0.005

On the lateral surface, LPMA activation increased concomitantly with visuospatial context, while OPA did not (ROI x Condition interaction: F(2,203)=14.7, p<0.0001; post-hoc ANOVAs – OPA: F(2,101)=0.38, p_corr_=0.99; LPMA: F(2,101)=10.48, p_corr_=0.0006; difference across conditions (i.e., difference of differences): LPMA versus OPA – Panorama versus Image: t(16)=2.771, p_corr_=0.0408, Cohen’s d=0.672; Panorama versus Street: t(16)=2.767, p_corr_=0.0411, d=0.671; Street versus Image: t(16)=5.21, p_corr_<0.001, d=1.26). The same pattern was present on the ventral surface – VPMA’s activation increased across the conditions, but PPA’s did not (ROI x Condition interaction F(2,203)=5.29, p=0.01; post-hoc ANOVAs – PPA: F(2,101)=3.58, p=0.078; LPMA: F(2,101)=8.19, p_corr_=0.0078; difference across conditions: VPMA versus PPA – Panorama versus Image: t(16)=0.13, p_corr_=0.99, Cohen’s d=0.078; Panorama versus Street: t(16)=2.68, p_corr_=0.048, d=0.065; Street versus Image: t(16)=2.83, p_corr_=0.036, d=0.69). On the medial surface, both the MPA and MPMA activity increased with visuospatial context, which likely reflects the high degree of overlap between these regions (Steel et al., 2021) (MPA: F(2,101)=12.95, p<0.001; post-hoc test: LPMA versus OPA – Panorama versus Image: t(16)=4.68, p_corr_<0.001, d=0.74; Panorama versus Street: t(16)=2.55, p_corr_=0.06, d=0.64; Street versus Image: t(16)=4.12, p_corr_<0.001, d=1.17; MPMA: F(2,101)=19.18, p<0.001; post-hoc test: Panorama versus Image: t(16)=5.61, p_corr_<0.001, d=0.88; Panorama versus Street: t(16)=3.15, p_corr_=0.018, d=0.84; Street versus Image: t(16)=5.02, p_corr_<0.001, d=1.51). However, consistent with the lateral and ventral surfaces, activity in MPMA was modulated by spatial context to a greater degree than MPA (ROI x Condition interaction: F(2,203)=4.24, p=0.023; post-hoc test: MPMA versus MPA – Panorama versus Image: t(16)=2.01, p_corr_=0.18, d=0.49; Panorama versus Street: t(16)=1.4, p_corr_=0.51, d=0.35; Street versus Image: t(16)=2.75, p_corr_=0.042, d=0.69).

To summarize, in contrast to the scene perception areas, all three place memory areas showed a strong effect of visuospatial context. Additionally, control regions occipital pole (V1) and fusiform face area (FFA) (Kanwisher et al., 1997), were not modulated by visuospatial context (V1: F(2,101)=1.68, p=0.20; FFA: F(2,50)=0.78, p=0.46; Figure 4). Further, activity of the hippocampus, which is involved in scene construction from memory (Spiers and Maguire, 2006; Hassabis and Maguire, 2009; Zeidman et al., 2015; Zeidman and Maguire, 2016; Barry et al., 2019) and represents the spatial scale of large environments (Baumann and Mattingley, 2013; Brunec et al., 2018; Peer et al., 2019) was not modulated by visuospatial context (F(2,101)=0.13, p=0.87; Figure 4). Together, these results emphasize the specificity of visuospatial context activity modulation to the place memory areas.

**Figure 4.**
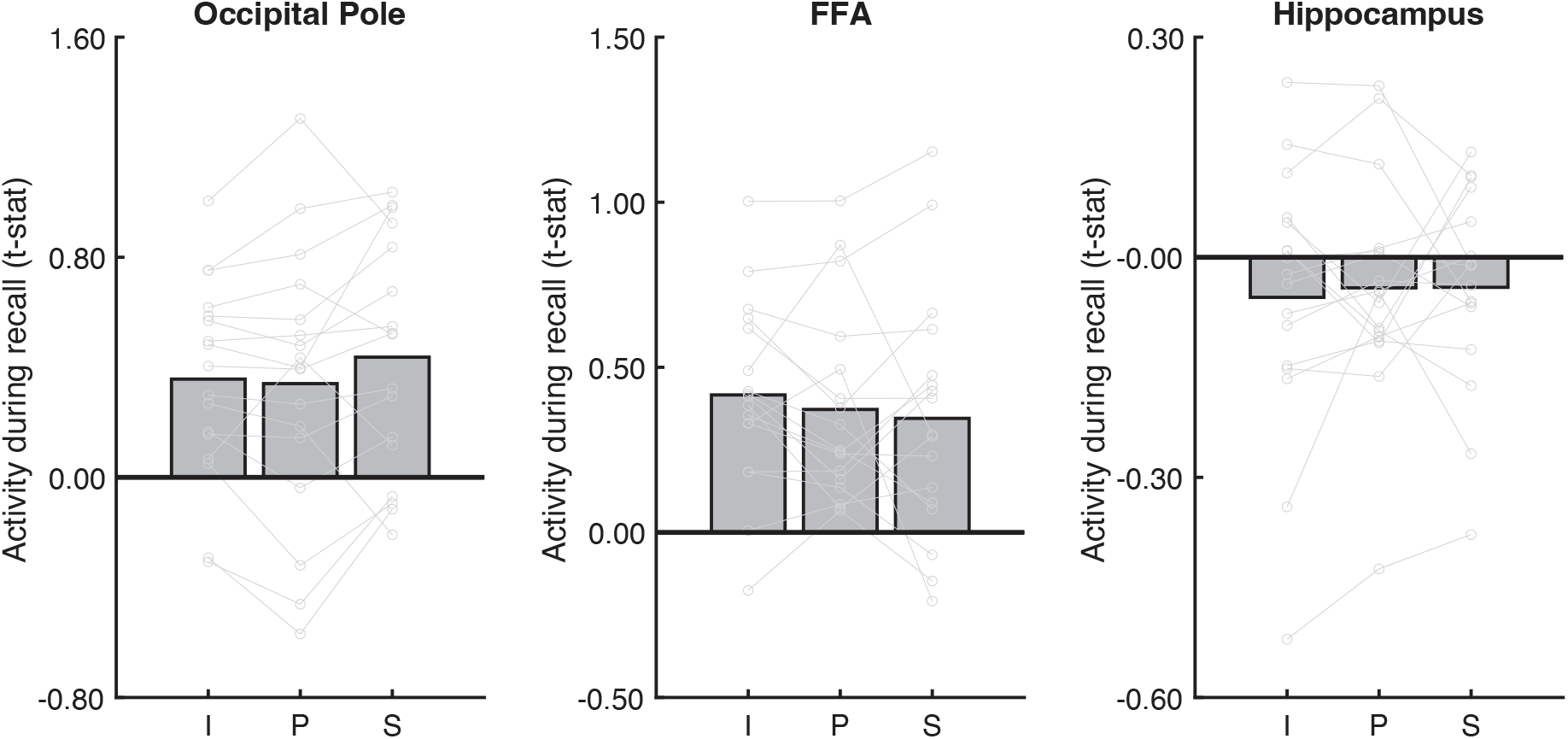
Analysis of control areas suggests that modulation by visuospatial context during recall is specific to the place memory areas. To determine whether visuospatial context was processed outside of the place memory areas, we considered additional regions for analysis: an early visual area (occipital pole, anatomically defined), a high-level visual area that selectively responds to faces rather than scenes (fusiform face area, FFA; functionally defined), and an area known to be involved in mnemonic processing (hippocampus, anatomically defined). No region was modulated by visuospatial context during recall. (I: Image, P: Panorama, S: Street)

In addition to exhibiting different activation patterns across the conditions, the place memory and scene perception areas showed markedly different overall activity magnitude during recall. Scene perception areas OPA and PPA were not activated above baseline in any condition (OPA – all ts<1.30, ps>0.21; PPA – all ts<1.11, ps>0.28). On the other hand, their respective place memory areas were significantly active in all conditions (LPMA – all ts>2.95, ps>0.009, ds>0.72; VPMA – all ts>6.99, ps>0.0001, ds>1.69). This pattern can be readily observed by examining each region’s activity timeseries (Figure 3B). This result, which is consistent with our prior work (Steel et al., 2021), suggests that the scene-selective perceptual areas may not play an active role in memory reinstatement, but instead are principally involved in processing visual input.

These univariate results suggest that the place memory areas process the extent of visuospatial context associated with a scene. But do the place memory areas actually represent the specific environment being recalled? To answer this question, we turned to multivariate analyses. We used an iterative split half correlation analysis to determine whether we could decode the recalled environment’s identity from the place memory areas. Importantly, to ensure that condition-related differences did not account for our ability to discriminate across the recalled environments, we calculated identity discrimination within each condition separately (i.e., only image vs image, panorama vs panarama, and street vs street discrimination was considered). We found that the place memory areas represented the stimulus identity during recall (LPMA: t(16)=3.13, p=0.0065, d=0.89; VPMA: t(16)=3.57, p=0.0026, d=0.95; MPMA: t(16)=4.71, p<0.001, d=1.14; Figure 5). Intriguingly, the scene perception areas also represented the scene being recalled (Lateral – OPA: t(16)=3.517, p=0.009, d=0.85; Ventral – PPA: t(16)=5.78, p<0.001, d=1.40; Medial – MPA: t(16)=4.558, p<0.01, d=1.11). On average, the scene perception areas contained significantly more identity-related information compared to our control areas V1, FFA, and hippocampus (perception areas vs V1: t(16)=2.77, p=0.012, d=0.67; vs FFA: t(16)=2.55, p=0.02, d=0.62; hippocampus: t(16)=4.93, p<0.0001, d=1.20), and the place-memory areas showed a similar trend (memory areas vs V1: t(16)=1.89, p_one-tailed_=0.0378, d=0.4607; vs FFA: t(16)=1.76, p_one-tailed_=0.043, d=0.4274; hippocampus: t(16)=4.36, p=0.0005, d=1.06). Because the OPA and PPA showed no univariate modulation during recall, we suggest that the stimulus-relevant information in these areas arises via feedback from the place memory areas.

**Figure 5.**
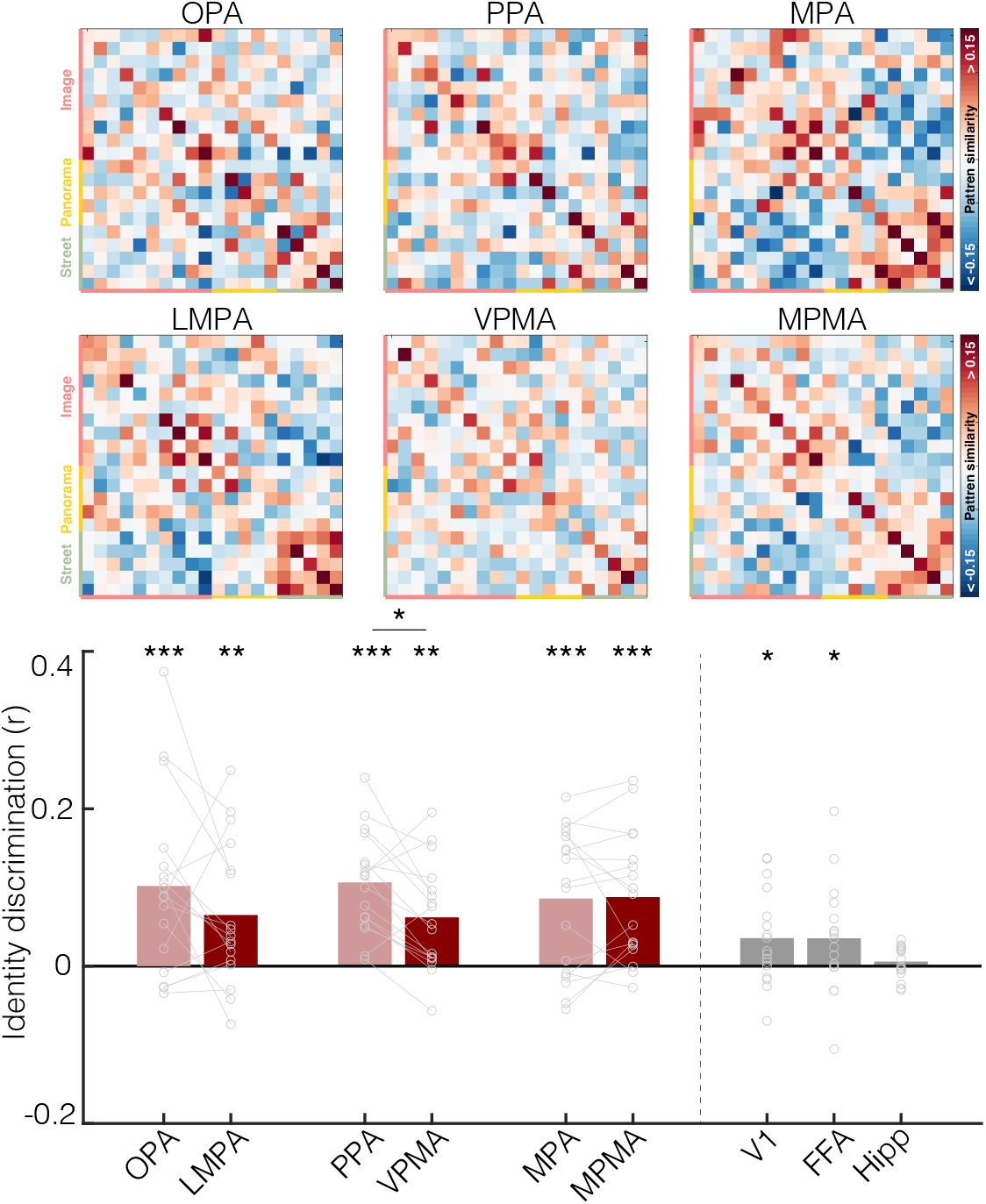
The place memory and scene perception areas represent the identity of the stimulus being recalled. Upper. Representational similarity matrix showing stimulus x stimulus correlation of the pattern of activity within each ROI. Each cell represents the similarity of activation patterns between two stimuli derived from an iterative half-split multi-voxel pattern analysis. The matrix diagonals represent the average similarity of a stimulus with itself across each data split (identity). Stimuli are grouped by condition (colored axis labels: image, panorama, street). Lower. Identity discrimination index for the place memory and scene perception areas. To ensure that condition-related differences could not explain identity decoding, we examined the ability to discriminate stimulus identity by comparing the average similarity between a stimulus and itself versus other stimuli within its condition. Both the scene perception and place memory areas evidenced significant identity discrimination (*=p<0.05, **=p<0.01, ***=p<0.005).

Together, the results of Exp. 1 confirmed that the place memory areas’ activity: i) reflects the extent of visuospatial context associated with the key scene view and ii) contains stimulus-specific information, as would be required if they represent the remembered local visuospatial environment.

### Place memory area activity reflects visuospatial context during visual recognition

Investigating neural activity during memory recall (Exp. 1) allowed us to look directly for regions processing unseen visuospatial information. Yet focusing on an explicit recall task has limitations. First, hypothetically, acting effectively in the real-world requires representing visuospatial context online along with incoming visual information (Haskins et al., 2020; Ryan and Shen, 2020; Rust and Palmer, 2021; Draschkow et al., 2022). Thus, we would expect a region that represents visuospatial context should also be involved in perception by implicitly reactivating contextual information from memory during recognition of perceived scene views. Second, methodologically, we do not know the precise content of participants’ imaginations, so magnitude differences in our explicit recall task could arise from the amount of information being recalled if participants recalled multiple scene views on each trial, or different scene views in different trials. Furthermore, although participants were instructed to recall a single field of view on each trial, internal reorientation during recall could activate the cortical head-direction network (Taube, 2007; Baumann and Mattingley, 2010; Shine et al., 2016; Nau et al., 2020) and potentially drive the apparent effect of visuospatial context observed in the place memory areas.

To address these limitations, we next turned to a visual recognition paradigm (Exp. 2). In Exp. 2, we briefly (500ms) presented the key scene view from each learned environment and participants performed an old/new judgment (Figure 6). We presented stimuli briefly to discourage participants from explicitly recalling unseen details and to ensure that the task primarily engaged recognition. In addition, we chose a recognition task to ensure that participants were not performing any mental reorientation that could activate the head direction network. Therefore, any difference in activation across the conditions would likely reflect implicit reactivation of visuospatial context rather than explicit, constructive mnemonic processes. We hypothesized that activity in the place memory areas would reflect the amount of known spatial context when performing a recognition task on a visually recognized scene. On the other hand, the scene perception areas would be active during the recognition task (because scenes are being perceived), but their activation magnitude would not reflect the amount of spatial context associated with the perceived scene.

**Figure 6.**
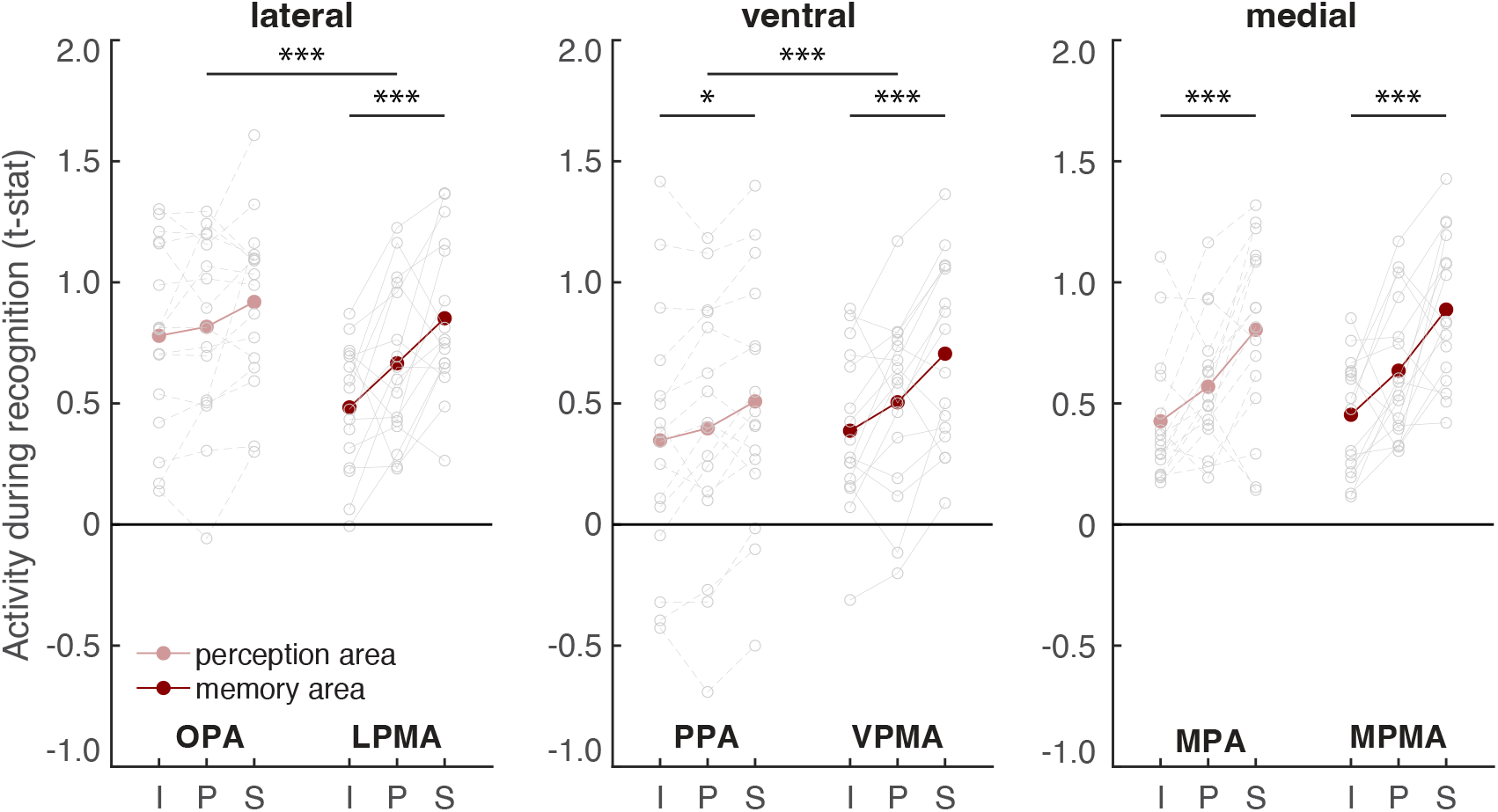
The extent of visuospatial context associated with a scene view modulates place memory area activation during the perceptual recognition task. A. We compared activation of the scene perception and place memory areas across visuospatial context conditions during recognition using an ANOVA with ROI (perception/memory area), visuospatial context (image, panorama, street [i,p,s]) and hemisphere as factors. On the lateral and ventral surfaces, the place memory area activation was modulated by visuospatial context to a greater degree than the scene perception areas. On the medial surface, activity of both the scene perception and place memory area was modulated by known visuospatial context. *: p<0.05, **: p<0.01, ***: p<0.005

Do the place memory areas process visuospatial context during recognition? Consistent with their roles in visual scene processing, all scene perception and place memory areas responded significantly during the recognition task (all ts>2.64, ps < 0.0187, ds>0.65). Importantly, however, we again found that visuospatial context specifically modulated activity in the place memory areas. On the lateral surface, visuospatial context impacted LPMA activity significantly more than OPA (ROI x Condition interaction: F(2,191)=6.95, p=0.003; post-hoc ANOVAs – OPA: F(2,95)=3.51, p=0.085; LPMA: F(2,95)=11.93, p=0.0004; difference across conditions: LPMA versus OPA – Panorama versus Image: t(15)=2.41, p_corr_=0.084, d=0.60; Panorama versus Street: t(15)=1.35, p_corr_=0.594, d=0.34; Street versus Image: t(15)=3.567, p_corr_=0.006, d=0.89). On the ventral surface, both PPA and VPMA activation was modulated by the amount of known visuospatial context. However, VPMA was modulated to a significantly greater degree (F(2,191)=9.95, p=0.005; post-hoc ANOVAs – PPA: F(2,95)=6.19, p=0.01; VPMA: F(2,95)=13.23, p=0.0002; difference across conditions: VPMA versus PPA – Panorama versus Image: t(15)=2.55, p_corr_=0.06, d=0.63; Panorama versus Street: t(15)=2.21, p_corr_=0.129, d=0.55; Street versus Image: t(15)=4.19, p_corr_<0.001, d=1.05). On the medial surface, both MPA and MPMA activation reflected visuospatial context, and these regions did not differ in their scaling across conditions (ROI x Condition – F(2,191)=1.75, p=0.19; Main effect of Condition – MPA: F(2,95)=12.94, p < 0.001; post-hoc test: Panorama versus Image: t(15)=2.18, p_corr_=0.135, d=0.746; Panorama versus Street: t(15)=3.15, p_corr_=0.018, d=1.01; Street versus Image: t(15)=4.50, p_corr_=0.0012, d=1.62; MPMA: F(2,95)=14.31, p < 0.001; post-hoc test: Panorama versus Image: t(15)=2.385, p_corr_=0.123, d=0.98; Panorama versus Street: t(15)=2.92, p_corr_=0.033, d=1.20; Street versus Image: t(15)=5.32, p_corr_<0.0001, d=2.24). Importantly, control ROIs FFA and V1 were not modulated by spatial context (V1: F(2,95)=0.29, p=0.74; FFA: F(2,47)=3.02, p=0.06; Figure 7). Although FFA approached significance, this was driven by lower activation to street images, which is opposite to the place memory areas. Likewise, hippocampal activity was not significantly modulated by visuospatial context during recognition (F(2,95)=3.02, p=0.06; Figure 7). While the effect of visuospatial context during recognition approached significance, this was driven by lower activation to panoramas compared to streets and images.

**Figure 7.**
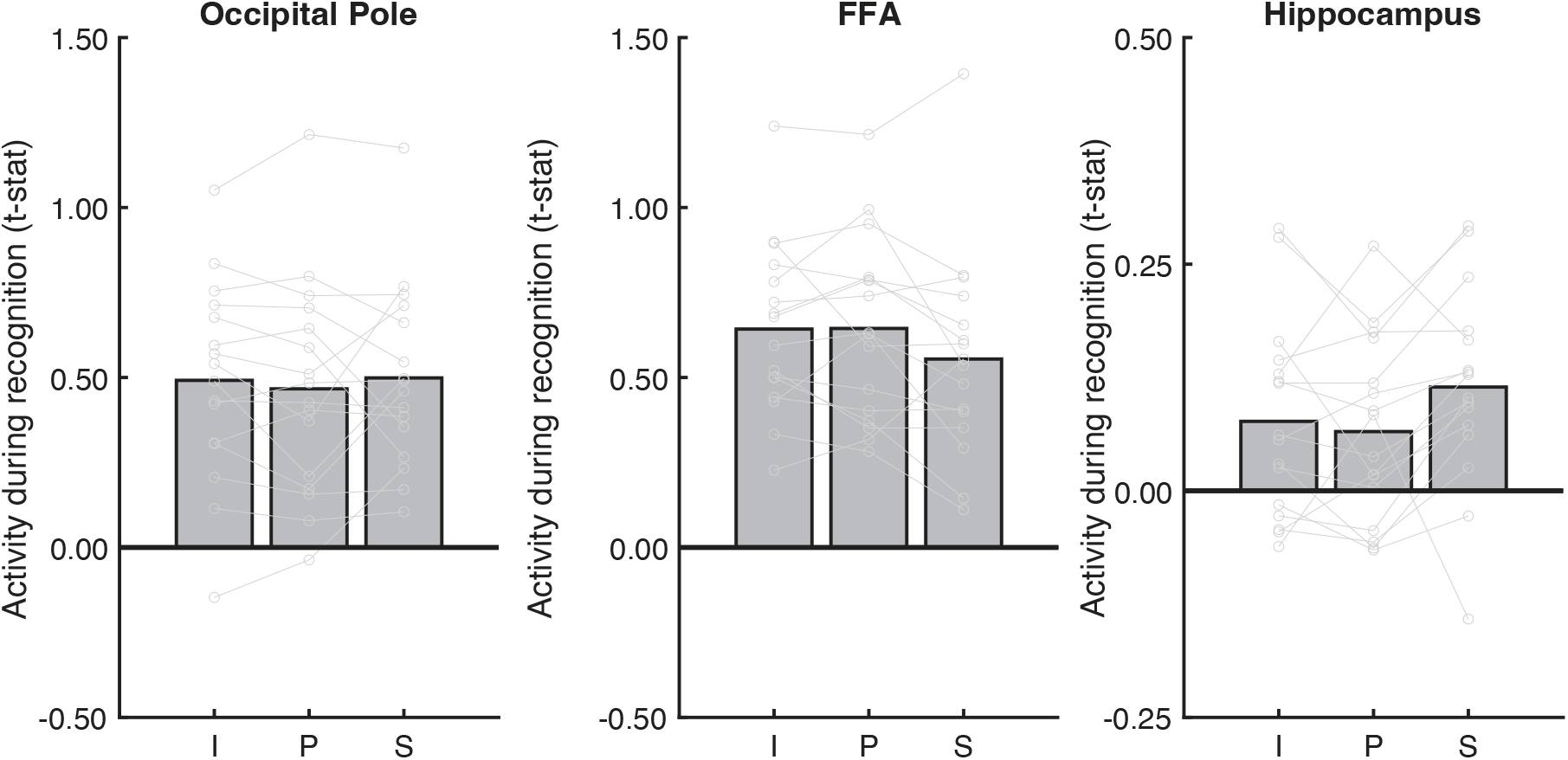
Analysis of control areas suggests that modulation by visuospatial context during recall is specific to the place memory areas. To determine whether visuospatial context was processed outside of the place memory areas, we considered additional regions for analysis: an early visual area (occipital pole, anatomically defined), a high-level visual area that selectively responds to faces rather than scenes (fusiform face area, FFA; functionally defined), and an area known to be involved in mnemonic processing (hippocampus, anatomically defined). No regions outside the place memory areas were significantly modulated by visuospatial context during recognition. I: image, P: panorama, S: Street

Finally, we wanted to determine whether the modulation by known visuospatial context was specific to the place memory areas. To answer this question, we turned to a whole-brain analysis (Figure 8). As we found in Experiment 1, the modulation by visuospatial context was remarkably specific: within posterior cerebral cortex, significant vertices fell largely within the place memory areas and did not appear in other category-selective areas. Notably, at the whole brain level, the lateral place memory area contained only a small number of significant vertices compared to the ventral and medial place memory areas; this is likely due to the heterogeneity in the location of the lateral place memory area across participants. Thus, the effect of visuospatial context on activation during recognition is largely restricted to the place memory areas.

**Figure 8.**
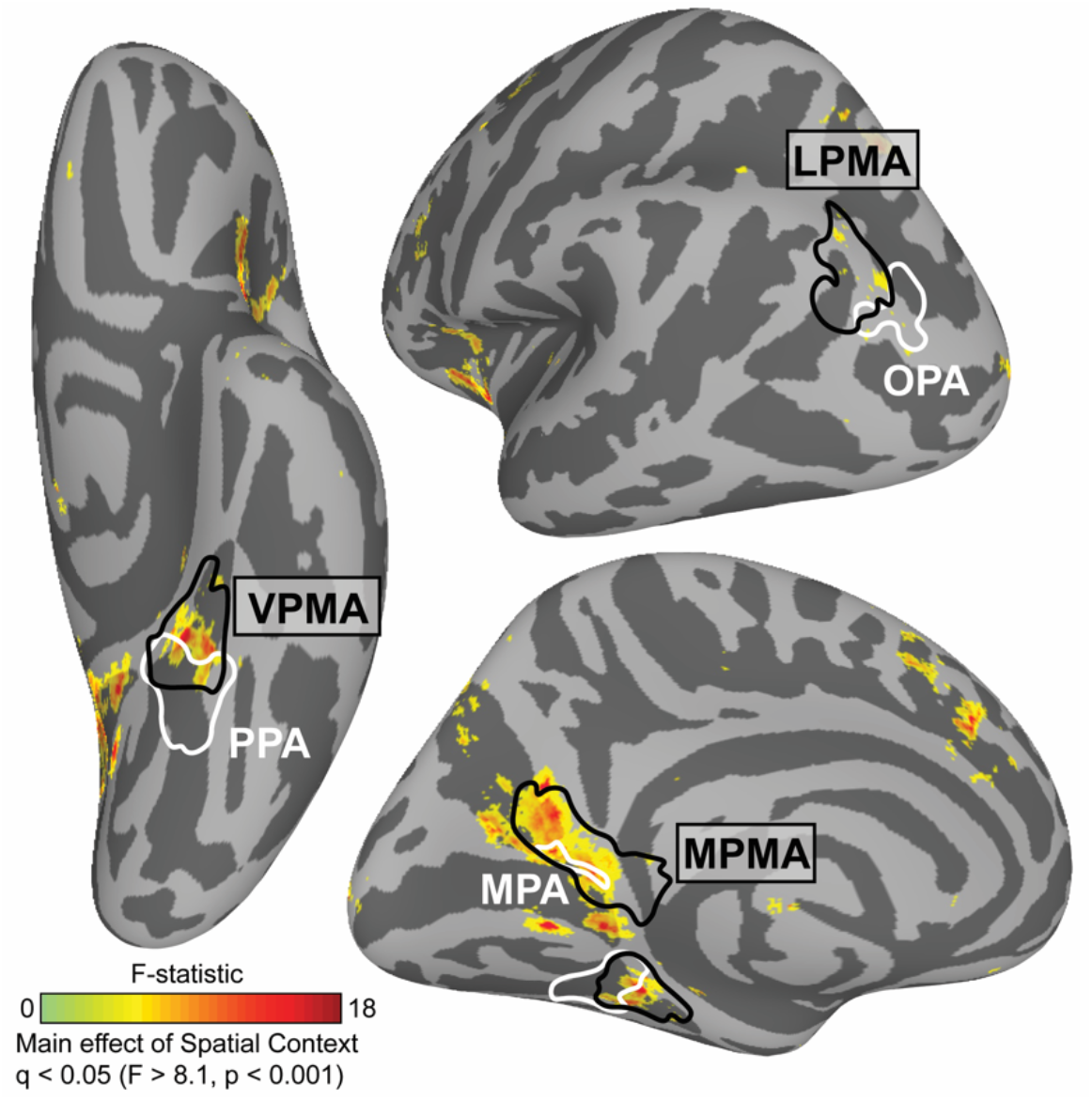
Modulation by spatial context across the brain during recognition. We compared neural activation across the visuospatial context conditions (Image, Panorama, and Street) using a vertex-wise ANOVA. We found that three clusters showed significant modulation by known spatial context, which fell largely within the place memory areas (black). Whole brain analysis thresholded at FDR-corrected p-value (q) < 0.05 (vertex-wise F>8.1, p<0.001; q < 0.05). Scene perception areas (white) and place memory areas (black) were defined using data from an independent group of participants (Steel et al., 2021).

The results of Exp. 2 suggest that, during visual recognition, the lateral and ventral place memory areas represent the visuospatial context associated with a visual scene to a greater degree than their perceptual counterparts, while MPA and MPMA both appear to process visuospatial context. Together with the result of Exp. 1, these data suggest that perceptual information from the visual system and contextual information from memory converge anterior to each of the scene perception areas, in the place memory areas.

### Post-scan behavioral results confirmed memory success

To ensure that systematic differences in i) subjective vividness and ii) memory strength could not explain the difference in neural activity across conditions, we had participants perform two behavioral assessments after MRI scanning. First, participants rated their ability to vividly imagine each stimulus on a 1-5 scale, with 5 being the most vivid. Participants reported strong overall ability to imagine the learned scenes (4.31 ± 0.5 STD) and reported no difference in imagery ability across the conditions (F(2,50)=3.03, p=0.06). Although this analysis approached significance, it reflects a decrease in vividness in the street condition compared to the image and panorama condition, which is in the opposite direction to our magnitude analysis results. Second, a subset of participants completed a naming test. In the naming test, participants were shown each key scene view and provided the name of the associated environment. Participants’ responses were highly accurate on this difficult memory assessment (0.86 ± 0.20 STD), and their accuracy did not differ across the conditions (F(2,32)=2.48, p=0.109). These results suggests that the observed differences in activation across conditions in the place memory areas reflects visuospatial context rather than other behavioral factors.

## Discussion

A central debate in neuroscience concerns the neural mechanisms facilitating the interplay of mnemonic and sensory information in the brain. This topic is particularly relevant for real-world scene perception, which requires representing properties of the environment that are outside the current field of view to inform ongoing perception and behavior (Robertson et al., 2016; Haskins et al., 2020; Berens et al., 2021; Draschkow et al., 2022). Yet, while the neural systems representing visual scenes are relatively well-understood (Epstein and Kanwisher, 1998; Maguire et al., 1998; Hasson et al., 2002; Malach et al., 2002; Epstein et al., 2007a; Dilks et al., 2013, 2022; Silson et al., 2015; Kamps et al., 2016; Malcolm et al., 2016; Bonner and Epstein, 2017a; Epstein and Baker, 2019; Lescroart and Gallant, 2019), how memory for the surrounding environment interfaces with scene processing is largely unknown. On one hand, visual cortex could jointly represent mnemonic and sensory information to facilitate memory-based perception (D’Esposito et al., 1997; O’Craven and Kanwisher, 2000; Rademaker et al., 2019). On the other hand, dedicated systems might respectively harbor mnemonic and sensory information to prevent mnemonic-perceptual interference (Spreng et al., 2009; Ranganath and Ritchey, 2012; Ritchey et al., 2015; Barnett et al., 2020; Ritchey and Cooper, 2020).

Our results suggest an intriguing third possibility. Here, we demonstrate that recalling a visuospatial environment activates a functional network immediately anterior and adjacent to the three scene-selective areas of high-level visual cortex, the “place memory network” (Steel et al., 2021). This arrangement could allow efficient mnemonic-perceptual interaction while minimizing mnemonic-perceptual interference. Although we previously identified the place memory network and described its responses to familiar stimuli and its connectivity, the nature of mnemonic processing in these areas was not clear (Steel et al., 2021). The present study offers three insights into the place memory areas’ mnemonic representations. First, by using virtual reality, we show that visuospatial information alone may be sufficient to drive activity in the place memory areas, implying that the episodic and multisensory details present in personally familiar memories may not be required. Second, we observed that the amount of known visuospatial context associated with a scene modulates the anterior areas’ activity during recall and recognition, suggesting that these areas process visuospatial content associated with a scene view. Third, using MVPA, we show that the place memory areas represent the specific environment being recalled, which we had not demonstrated in our prior work (Steel et al., 2021). Taken together with prior work showing an anterior shift for mnemonic compared with visual information for scenes (Baldassano et al., 2013, 2016; Rugg and Thompson-Schill, 2013; Silson et al., 2016, 2019; Peer et al., 2019; Bainbridge et al., 2020; Steel et al., 2021; Srokova et al., 2022), these results suggest that the place memory areas likely play a key role in memory-guided scene perception by processing the remembered visuospatial context of a scene.

The conventional view of scene perception is that integration of mnemonic and perceptual information occurs in the MPA (a.k.a., retrosplenial complex) (Malcolm et al., 2016; Epstein and Baker, 2019; Dilks et al., 2022). This view is based on reports showing that mnemonic tasks engage MPA. For example, unlike OPA and PPA, MPA responds preferentially to familiar versus unfamiliar images, and MPA activates more than OPA or PPA in memory tasks that involve linking a scene view with its environment context (Epstein et al., 2007b). Further, unlike OPA and PPA, MPA jointly represents both egocentric viewpoint of the current scene and the viewer’s allocentric position within the environment (Vass and Epstein, 2013; Marchette et al., 2014; Shine et al., 2016; Nau et al., 2020). In contrast, PPA and OPA are implicated in perceptually-oriented functions: scene categorization (Persichetti and Dilks, 2019; Dilks et al., 2022) and visually guided navigation (Julian et al., 2016; Bonner and Epstein, 2017b; Lescroart and Gallant, 2019), respectively (Dilks et al., 2022). Thus, the scene perception areas appeared to differ in their degree of mnemonic function.

Our results significantly broaden our understanding of how mnemonic and perceptual information interface during scene processing. Although the amount of known visuospatial context largely did not impact PPA and OPA activation (consistent with the conventional view), we found that activation of the cortical territory immediately anterior to each of these regions is modulated by remembered visuospatial context. Thus, while the function of OPA and PPA may be more directly linked to perception, mnemonic information likely influences representations in these regions via their anterior mnemonic counterparts. Consistent with this view, we found that OPA and PPA represent the identity of the recalled scene, which we hypothesize arises via feedback from the place memory areas. Therefore, memory information likely plays a larger role in the overall function of OPA and PPA than the conventional view of scene processing suggests. A second, related hypothesis concerns the rigidity of visual representations in the scene perception versus place memory areas. Specifically, it could be that the scene perception areas focus only on the present image (i.e., the scope of information processed by these areas is limited to the current field of view); In contrast, place memory areas may have more malleable fields of view, such that the amount of information they represent may depend both on the extent of memory for one’s surroundings and the scope of one’s internal attention based on current task-demands (i.e., are you remembering the visuospatial context around the immediate corner, or planning a longer journey through a city). This could relate to the slight decrease in identity decoding within the place memory areas compared to the scene perception areas during recall if the amount of visuospatial information within the place memory areas is not fixed across trials. This may also relate to boundary transformations like extension and contraction (Intraub and Richardson, 1989; Bainbridge and Baker, 2020; Hafri et al., 2022; Gandolfo et al., 2023). Thus, our results show a new network of brain areas processing mnemonic information regarding visuospatial context associated with a scene is interposed between structures implicated in scene memory (e.g., the hippocampus or MPA) and the perceptually-oriented scene areas (i.e., OPA and PPA), and future work should consider these alternative accounts of the memory area’s activity.

Our findings in MPA largely agree with the conventional understanding of this region’s function. Unlike the OPA and PPA, the degree of associated visuospatial context modulated MPA activation in both memory tasks. Prior work has implicated MPA in representing visuospatial context. For example, boundary extension, considered a proxy for representing visual information out of view, has been linked to MPA (Park et al., 2007). Likewise, panoramic content associated with a scene can be decoded from MPA (Robertson et al., 2016; Berens et al., 2021). In addition, MPA also represents world-centered (allocentric) facing direction (Vass and Epstein, 2013; Marchette et al., 2014; Julian et al., 2018), which requires knowledge of the environment’s spatial layout (Marchette et al., 2014). Activity in medial parietal cortex near MPA becomes more correlated with the hippocampus after learning a new environment, suggesting that this region is important for encoding new place memories (Steel et al., 2019). Further, damage to this area causes severe topographic agnosia while leaving visual recognition abilities intact (Takahashi et al., 1997; Aguirre and D’Esposito, 1999). Notably, unlike OPA and PPA, which lie immediately posterior to a place memory area, MPA is largely contained within its corresponding place-memory area (i.e., MPMA (Steel et al., 2021)). Together, our findings support the conclusion that MPA could be considered a visually-responsive sub-region of a largely mnemonic cortical area and may be different in kind from PPA and OPA, which play an active role in visual analysis.

More broadly, our findings raise important questions about how perceptual and mnemonic neural signals converge. Several studies have noted anterior shifts in activation when people visualize versus perceive stimuli (Rugg and Thompson-Schill, 2013; Favila et al., 2018, 2020; Silson et al., 2019; Bainbridge et al., 2020; Popham et al., 2021; Steel et al., 2021; Srokova et al., 2022). What representational transformation underlies this shift? One account posits that this shift reflects a transformation from sensory information (i.e., the visual properties of a house) in high-level visual areas to more abstract, amodal, semantic representations (i.e., the mental concept of a house) (Chao et al., 1999; Huth et al., 2012; Margulies et al., 2016; Huntenburg et al., 2018; Popham et al., 2021). For example, one recent study compared a computational language model fit to movie watching versus listening to stories and concluded that the anterior shift reflected a transformation from visual features to representations grounded in language (Popham et al., 2021). However, listening to stories engages many processes, including top-down imagery, working memory, and narrative understanding, so pinpointing the nature of these representations using naturalistic paradigms is challenging.

In contrast, our results argue against a purely semantics-based account of the anterior shift. By focusing on one visual process (visual scene analysis) and isolating visuospatial context, we found that the anteriorly-shifted activity reflects visuospatial context currently out of sight. As such, our findings are not well accounted for by language-based accounts of the anterior shift. Instead, we hypothesize that these anterior areas contain visuospatial information relevant to making memory-based predictions of the visuospatial environment in service of perception (Steel et al., 2023). Notably, because we focused only on visuospatial context, we cannot say whether the anteriorly-shifted representations are exclusively visuospatial. One limitation of our approach (i.e., modulating the amount of visuospatial content) is that increasing visuospatial context necessarily entails the addition of other information as well as increased ecological naturalism. For example, visual information on the left and right side of the street could include details like houses or shops that provide semantic details about the environment, and this information would be more prevalent in the panorama and street conditions (versus the image condition). Relatedly, the presence of the occluder in the image condition diminishes its naturalistic and immersive quality compared to the panorama and street conditions. Therefore, while our results demonstrate that episodic and multisensory experiences are not necessary to elicit the anterior shift for mnemonic information, we cannot rule out that other information could elicit this anterior shift, such as semantic associations, multi-sensory context, and motor memories related to head movements made during stimulus encoding, and the feeling of immersion. Could anteriorly-shifted representations even reflect amodal, semantic-based information (Popham et al., 2021)? Future studies will be needed to understand the nature of the representational transformation anterior to visually-responsive cortex, and how these principles generalize outside the case of scenes.

## Conclusion

In conclusion, by using VR to precisely manipulate the visuospatial context associated with a scene view, we found that perceptual representation of the visual scene in front of us and our memory of the surrounding environment converge at the anterior edge of visually-responsive cortex in the place memory areas. This functional architecture could allow efficient interaction between the representation of the current visual scene with our memory of the wider visuospatial context.

## Acknowledgments

The authors would like to thank Yeo Bi Choi and Terry Sackett for assistance with data collection. This work was supported by grants from the National Science Foundation (#2144700) and NVIDIA to CER. AS is supported by the Neukom Institute for Computational Science.

## Notes

### Competing Interest Statement

The authors have declared no competing interest.

### Summary of Updates

update with additional analyses, figures, and main text

